# ECLIPSE: Exploration of Complex Ligand-Protein Interactions through Learning from Systems-level Heterogeneous Biomedical Knowledge Graphs

**DOI:** 10.1101/2025.11.05.686382

**Authors:** Heval Atas Guvenilir, Tunca Doğan

**Author notes:** Current affiliation of the author: Department of Bioinformatics, Fraunhofer Institute for Algorithms and Scientific Computing (SCAI), 53757 Sankt Augustin, Germany.

## Abstract

Discovering new, efficacious molecules remains slow and costly; rigorous data science-driven systems-level approaches are therefore essential to prioritise hypotheses and de-risk drug development. In this study, we present ECLIPSE, a systems-level framework for compound/ligand–protein interaction (CPI) representation and prediction, combining heterogeneous knowledge graphs (KGs), which encode large-scale entity–relation structure, with graph neural networks that exploit relational inductive biases to perform inference on graph-structured data. ECLIPSE uses our comprehensive biomedical KG-based platform, CROssBAR, incorporating genes/proteins, drugs, compounds, pathways, diseases, and phenotypes, along with their multi-layered relationships. Each entity is assigned input features derived from language or graph representation learning models and projected via type-specific neural network layers. To process these featurized biomedical KGs for bioactivity prediction, we employed the heterogeneous graph transformer (HGT) architecture. In contrast to the majority of GNN algorithms, which are restricted to homogenous graphs, HGT can handle graph heterogeneity and maintain node-and edge-type dependent representations through its attention mechanism. ECLIPSE achieves strong performance on challenging, protein-family– specific CPI benchmarks compared with baseline and state-of-the-art methods; ablations confirm performance gains from modelling graph heterogeneity and all feature sources. Use-case analyses on a druggable kinase (PIM1) and a historically undruggable receptor (HER3) illustrate generalizability across target classes and activity ranges. By leveraging direct and indirect relationships embedded in biomedical KGs, ECLIPSE provides context-aware CPI inference that is scalable to real-world settings. Code, datasets, and trained models are released to support reproducibility and reuse.

## 1. Introduction

Drug discovery is a complex process involving the identification and optimisation of compounds that interact selectively with intended target biomolecules, aiming to produce the desired therapeutic effect. In recent years, computational approaches have become essential across various stages of the drug discovery and development pipeline ^1–8^. Among these applications, virtual screening plays a central role in early-phase drug discovery by enabling the rapid identification of promising drug candidates from extensive compound libraries prior to experimental procedures. The remarkable ability of artificial intelligence (AI) based analytical methods for large-scale data handling and pattern discovery has led to their widespread adoption in this process^9^.

Both traditional machine learning (ML) ^10–13^ and cutting-edge deep learning (DL) algorithms ^14–17^ have been used for virtual screening. Among these, CGKronRLS is a model that employs a regularised least squares (RLS) algorithm with a Kronecker product kernel, combining drug and target similarities for bioactivity prediction ^11^. DeepDTA ^14^ and MDeePred ^15^ are popular DL models developed for predicting compound-protein binding affinity. DeepDTA employs two stacked CNN blocks to process drug SMILES and protein sequences separately. MDeePred utilises a pairwise input neural network (PINN) architecture comprising a protein input CNN and a compound input feed-forward neural network. The method incorporates multiple types of protein features, including sequence, structural, evolutionary, and physicochemical properties as 2-D vectors, and uses ECFP4 fingerprints for compounds. Although current ML/DL methods show promise in predicting compound-protein interactions (CPIs)/drug-target interactions (DTIs), the dynamic and complex nature of biological systems introduces numerous factors— such as off-target effects, disease heterogeneity, cellular uptake, and drug metabolism—that ultimately affect the outcome. Therefore, relying solely on basic virtual screening approaches is inadequate to address this biological problem. Conversely, adopting a systems-level strategy that integrates direct and indirect relationships in molecular and cellular processes, including protein-protein interactions and signaling/metabolic pathways, together with high-level concepts such as protein-disease relationships, drug-disease indications, pathway-disease modulations, and phenotypic implications, can increase the likelihood of success in drug discovery. ^18–21^.

Advancements in data analysis techniques have facilitated the processing and interpretation of large-scale biomedical data. One of the most promising relationship-centric data formats for this purpose is the knowledge graph (KG), which can represent complex associations between different layers of biomedical data. Knowledge graphs encode information in a network format, where nodes represent entities and edges represent relationships or interactions between them. One common approach for constructing biomedical KGs is extracting information from unstructured text in biomedical literature, often from databases like PubMed via text mining, which are then represented as subject-predicate-object semantic triples ^22–24^. Another emerging strategy involves integrating diverse data types from multiple structured sources. This approach relies on cross-referencing between these sources to map and consolidate information, resulting in comprehensive KGs enriched with varied biomedical data. Successful applications of these integrative approaches include HetioNet ^25^, BioGrakn ^26^, CROssBAR ^18^, Bioteque ^19^, SPOKE ^21^, and OREGANO ^20^.

Given the extensive and multifaceted nature of these biomedical KGs, it is crucial to develop robust methods that can learn from their complex structure to predict biological relationships accurately. In this context, graph representation learning (GRL) techniques become indispensable. GRL refers to the process of learning low-dimensional representations (i.e., embeddings) of graphs and their components while preserving structural and relational characteristics. Shallow embedding methods like matrix factorization (e.g., Laplacian eigenmaps) and random walk (e.g., Node2Vec, DeepWalk), along with basic KG embedding methods (e.g., TransE, RESCAL), are commonly employed GRL algorithms for the extraction of meaningful features from graph-structured data. These features can be utilized across various downstream ML tasks or exploratory data analyses ^27,28^. With the advancement of DL approaches, graph neural networks (GNNs) have emerged as a promising modelling technique within the domain of GRL. GNNs operate by iteratively updating node representations based on information propagated from neighboring nodes in the graph. This iterative mechanism enables GNNs to capture both local and global graph structures, thereby facilitating the acquisition of rich node (or edge) representations.

In the context of CPI prediction, the predominant usage of GRL algorithms involves treating each protein and/or drug/compound structure as an individual graph. In this representation, atoms/amino acids are depicted as nodes, and chemical bonds connecting these entities are represented as edges ^16,17,30^. While successful in capturing the structural characteristics of proteins and drugs/compounds, these approaches remain limited to basic virtual screening. Newer applications that integrate relational biomedical data are primarily focused on drug-drug, protein-protein similarity/interaction networks and/or protein-drug interaction bipartite graphs ^31–34^. Although some efforts have emerged to benefit from additional types of biomedical data, including disease and side effect associations of proteins and drugs ^35–38^, these applications typically rely on similarity-based calculations and shallow graph embedding techniques, often followed by traditional ML/DL classifiers/regressors or simple encoder/decoder systems. Overall, the predominant reliance of traditional CPI prediction methods on structural and molecular similarities between proteins and compounds limits their ability to generalize beyond known interaction patterns. These approaches fall short of adopting system-level approaches that harness comprehensive heterogeneous biomedical data.

In this study, we introduce ECLIPSE, a novel framework for representing and predicting CPIs at a systems level. ECLIPSE addresses the generalisation-related problems of conventional ML/DL-based CPI prediction methods by adopting a graph-based DL framework that captures multi-layered, semantically related heterogeneous biomedical relationships. Serving as a hybrid approach, it incorporates learned embeddings of various biological entities into heterogeneous graph data for better representation and more effective processing of complex relationships.

Our approach aims to uncover deeper relationships between proteins and compounds, leading to more reliable CPI predictions applicable in real-world scenarios. As its input data, ECLIPSE utilises large-scale biomedical KGs constructed by CROssBAR ^18^. CROssBAR is a comprehensive system that integrates a wide range of publicly available and reliable biomedical data sources including UniProt ^39^, IntAct ^40^, ChEMBL ^41^, DrugBank ^42^, Reactome ^43^, KEGG ^44^, OMIM ^45^, and Human Phenotype Ontology ^46^ to build computable heterogeneous KGs and to provide a broad spectrum of biological information. CROssBAR KGs present sub-networks of interconnected entities including genes/proteins, biological processes/pathways, diseases, phenotypes, drugs and compounds. These entities are linked through a variety of relationships such as protein-protein interactions, protein-pathway associations, drug-disease indications, and disease-phenotype associations. ECLIPSE employs the heterogeneous graph transformer (HGT) architecture ^47^ to learn from CROssBAR KGs, and incorporates initial node embeddings derived from various pretrained representation models ^48,49^. Through the attention mechanism of the HGT algorithm, ECLIPSE extracts node-and edge-type dependent representations, allowing it to capture the complex patterns of biological relationships.

To evaluate the performance of ECLIPSE, we carried out experiments on the ProtBENCH transferases bioactivity benchmark dataset ^50^, which includes diverse stratified data splits. We compared the results with those obtained from various baseline models and benchmarked ECLIPSE models against prominent CPI prediction methods from the literature. We also performed a data-centric ablation study to investigate the effect of different biological components and their relationships on learning CPIs. To demonstrate the model’s utility, we evaluated its capacity to predict values at the tails of the bioactivity range. Lastly, we conducted a use-case study based on the predictions for druggable and non-druggable protein samples to further assess the robustness and reliability of our system. The entire framework, including datasets, results, and models, is available as an open-access repository (https://github.com/HUBioDataLab/ECLIPSE), promoting reproducibility and encouraging collaboration within the scientific community.

## 2. Results and Discussion

This chapter comprises multiple subsections that present a comprehensive evaluation of the ECLIPSE system. The first subsection (2.1) provides a general summary regarding the proposed method. The second subsection (2.2) presents the performance evaluation of the ECLIPSE models, including a comparison against different baseline models. The third subsection (2.3) compares ECLIPSE models with state-of-the-art (SOTA) bioactivity prediction models on the well-known Davis kinase benchmark dataset ^51^. The fourth subsection (2.4) examines the impact of various graph node/edge types (i.e., biological components) on learning through an ablation study. In the fifth subsection (2.5), the applicability of the ECLIPSE system is examined for the prediction of bioactivity range tails. The final subsection (2.6) presents a use-case study based on predictions for druggable and undruggable proteins, further assessing the reliability and robustness of the ECLIPSE framework.

### 2.1. The Overview of ECLIPSE

ECLIPSE is a systems-level representation and prediction framework for bioactivity prediction of drug candidate compounds against human target proteins. Figure 1 displays the schematic workflow of the ECLIPSE system. It utilises CROssBAR’s integrated biomedical KG to extract complex relationships between biological entities. The process starts by sampling subgraphs from the integrated CROssBAR KG and projecting initial node features (see Methods 4.2) into fixed-size embeddings using type-specific MLPs. These embeddings are then fed into the HGT layers. Through heterogeneous mutual attention, heterogeneous message passing, and target-specific aggregation steps, the HGT algorithm embeds each target node by processing its own features and those of its neighbours. In the heterogeneous mutual attention step, attention vectors are obtained, which are used to weight the corresponding messages during the message passing process. After this, HGT aggregates the information from source nodes to target nodes through the target-specific aggregation step, resulting in updated vectors. This is followed by mapping the updated vector of each target node back to its type-specific distribution. A key benefit of employing HGT is its consideration of edge specificity, meaning that the algorithm computes distinct weight matrices for each edge type, allowing for the generation of contextualised representations for each target node. This capability effectively addresses graph heterogeneity. After learning node representations through the HGT module, the updated compound and protein embeddings are passed through separate MLPs for further refinement. Then, ECLIPSE performs an edge regression task for bioactivity modelling. The task involves either calculating the dot product of compound and protein embeddings or concatenating them and passing through fully-connected (FC) layered neural networks, depending on the selected model design. For model training and evaluation, ECLIPSE utilises the ProtBENCH transferases bioactivity benchmark dataset ^50^.

**Figure 1.**
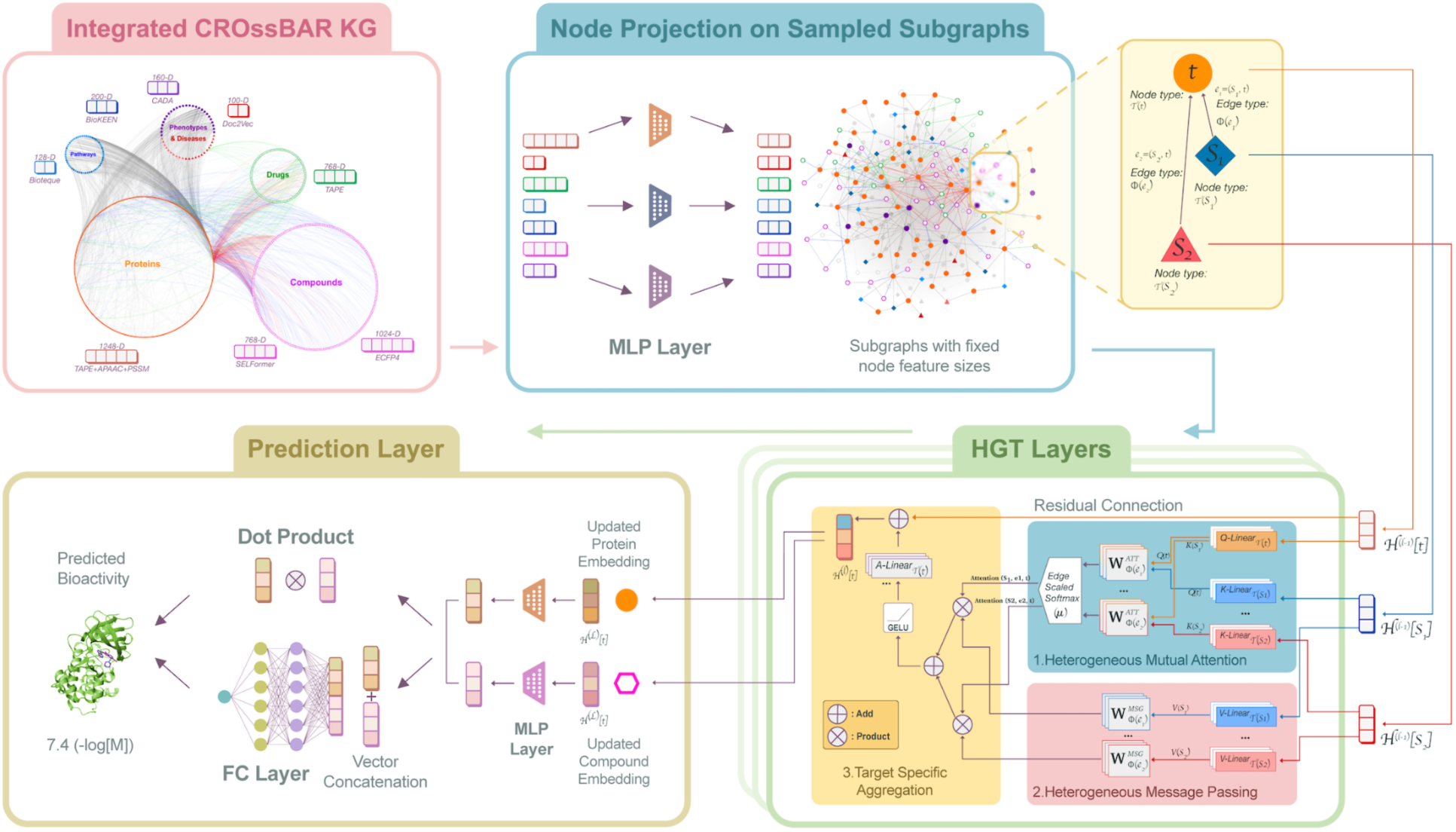
The schematic representation of the ECLIPSE framework for predicting compound– protein interactions. The **Integrated CROssBAR KG** module provides a multi-relational biomedical graph of proteins, compounds, drugs, pathways, phenotypes, and diseases, serving as the structural foundation for representation learning. In the **Node Projection on Sampled Subgraphs** module, type-specific MLP layers project heterogeneous input node features into fixed-size representations. These embeddings are then passed into stacked **HGT Layers**, which apply heterogeneous mutual attention, message passing, and target-specific aggregation with residual connections to generate contextualised node embeddings. Finally, the **Prediction Layer** combines updated compound and protein embeddings, which are first refined through separate MLPs, either through vector concatenation with a fully-connected network or via dot product, to predict bioactivity values.

### 2.2. Construction and Evaluation of Different ECLIPSE Models

We developed several versions of the ECLIPSE model, alongside baseline models, to provide a comprehensive assessment for the CPI prediction task. First, we built two alternative architectures, differing in the final prediction module (i.e., the way learned embeddings of protein and compound nodes are processed), one of which is based on dot-product (DP) computation and the other employs a fully-connected (FC) neural network. Subsequently, we constructed two versions of ECLIPSE regarding the input compound feature vectors: (i) models incorporate ECFP4 fingerprints, and (ii) SELFormer embeddings. ECFPs are commonly used circular topological fingerprints that capture the presence of particular substructures in a compound molecule ^52^. On the other hand, SELFormer is a transformer-based chemical language model (CLM) that utilises SELFIES notations of compounds to generate molecular embeddings ^49^. Overall, four distinct models were obtained: ECLIPSE-DP_SELFormer, ECLIPSE-DP_ECFP4, ECLIPSE-FC_SELFormer, and ECLIPSE-FC_ECFP4 (see Methods 4.2 and 4.3).

Additionally, we constructed random forest (RF) regression models as baselines, since RF is a widely used and well-performing algorithm in CPI prediction. Baseline RF models utilised the same initial protein and compound node features of ECLIPSE models as input feature vectors to yield a fair comparison. We also developed baseline models derived from the ECIPSE architecture by removing the HGT module, allowing us to isolate and assess its contribution.

These baseline models processed the initial protein and compound embeddings either by computing their dot product after linearization (model names: DP_SELFormer and DP_ECFP4) or concatenating the linearised vectors of initial compound and protein features and then passing them through multiple FC layers (model names: FC_SELFormer and FC_ECFP4).

We trained and tested these models using transferases protein family dataset (a subset of our previously developed ProtBENCH family-specific large-scale CPI datasets from ^50^), which play a critical role in drug discovery. We trained and tested our models utilising three different train-test data splits: (i) “random-split” (for predicting known inhibitors for known targets, e.g., drug repurposing), (ii) “dissimilar-compound-split” (for predicting novel inhibitors for known targets), and (iii) “fully-dissimilar-split” (for predicting new inhibitors for new targets). We optimised the hyperparameters (Table S1) of our models using a random search, by splitting our training folds into train/validation sets with a 95/5 ratio. The total runtime of models ranged from 10 minutes to 5 hours, depending on the model architecture and selected hyperparameters. These computations were performed using an NVIDIA RTX A5000 graphics card equipped with 24 GB of GPU memory. The selected (optimal) hyperparameter sets for each split are provided in Table S2. Model test performance results are given in Figure 2 and Table S3.

**Figure 2.**
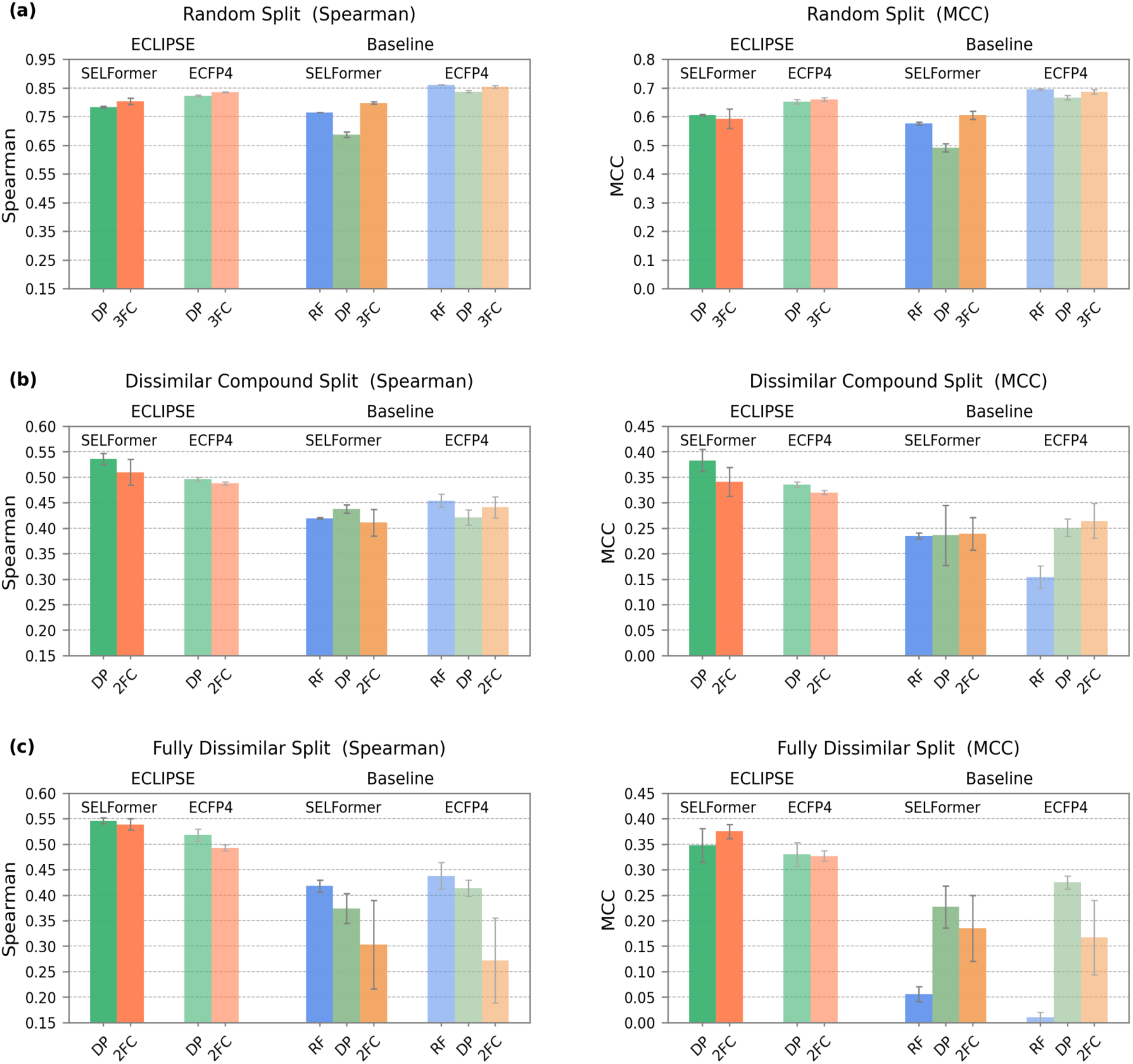
Bar plots of Spearman and MCC scores of ECLIPSE and baseline models on the **(a)** random, **(b)** dissimilar-compound, and **(c)** fully-dissimilar splits of the transferases test dataset (DP: dot product, RF: random forest, FC: fully-connected neural network).

In Figure 2, bar plots represent the test performance scores of the models, as measured by the Spearman correlation coefficient and the MCC metrics. Each model was trained and evaluated across five independent runs to derive error bars and assess performance variability. For statistical testing, we applied the Almost Stochastic Order (ASO) significance test using the deep-significance tool (https://github.com/Kaleidophon/deep-significance), which is specifically designed for DL models. Unlike traditional mean-comparison methods, the ASO test compares the entire distribution of scores and computes an epsilon minimum (eps_min) value. A model is considered to outperform another (reported as being stochastically dominant over) when its eps_min value is below 0.5 ^3^. Epsilon minimum values for all pairwise model comparisons (with a confidence level of 0.95) are provided in Supplementary Data Table 1, where Bonferroni correction was applied to control the family-wise error rate.

### ECLIPSE vs. Baselines

Across the challenging scenarios (i.e., the dissimilar-compound-split and fully-dissimilar-split), all ECLIPSE models consistently outperform the baseline models, demonstrating stochastic dominance with high statistical confidence based on ASO scores (eps_min = 0 for nearly all pairwise comparisons at 0.95 confidence level; see Figure 2b, 2c, and Supplementary Data Table 1). This robust performance highlights the advantage of incorporating heterogeneous graph information for CPI modeling. By effectively integrating diverse biological relationships, ECLIPSE models exhibit improved generalization capability and resilience against overfitting.

Although baseline models using ECFP4 fingerprints (i.e., RF_ECFP4, DP_ECFP4, and FC_ECFP4) achieve higher performance in the random-split setting (with eps_min values ranging from 0.005 to 0.551 at 0.95 confidence level; see Figure 2a and Supplementary Data Table 1), this outcome likely results from the simplified nature of random splits, where highly similar compounds are shared between training and test sets. Such conditions can artificially inflate performance metrics due to model memorization rather than true learning of generalizable features ^50^. In contrast, the strong performance of ECLIPSE models across more stringent splits confirms their reliability and suitability for realistic CPI prediction scenarios where novel and structurally diverse compounds are encountered.

### ECLIPSE Model Variants

When comparing the ECLIPSE models with each other, those based on SELFormer representations consistently exhibit superior or competitive performance relative to models relying on ECFP4 fingerprints in challenging scenarios, highlighting the effective representation capabilities of CLM-based molecular embeddings. Similarly, DP-based architectures generally achieve slightly better performance than FC-based models under these challenging conditions, likely due to their parameter efficiency and reduced complexity. However, this performance trend reverses in random-split scenarios, where ECFP4-based and FC models outperform their SELFormer-based and DP counterparts, respectively. This reversal likely occurs because random splits preserve chemical similarity between training and test sets, enabling ECFP4 fingerprints’ explicit substructural encoding and FC layers’ higher parameter capacity to capture familiar patterns effectively. Consequently, SELFormer and DP architectures excel in extrapolating to novel chemical spaces, whereas ECFP4 and FC models are superior in interpolation tasks. In summary, the remarkable performance of ECLIPSE models in challenging situations is promising and implies their strong adaptability to real-world CPI prediction tasks.

### 2.3. Performance Comparison with State-of-the-Art (SOTA) Models

For the comparison against SOTA CPI prediction models in the literature, we trained and tested ECLIPSE with the widely used Davis kinase benchmark dataset (see Methods 4.1 for details), and retrieved the results of SOTA models directly from the MDeePred ^15^ and MGraphDTA ^17^ studies. According to the results outlined in Table 1, ECLIPSE-FC_ECFP4 consistently outperformed SOTA models. It stands out as the top-performing model in all regression metrics, including CI, RMSE, and Spearman correlation, as well as in the MCC classification metric for both 1 μM and 100 nM bioactivity thresholds. Overall, ECLIPSE models with fully-connected layers performed better than their dot-product counterparts, and models incorporating ECFP4 fingerprints exhibited higher performance compared to those using SELFormer embeddings.

**Table 1.**
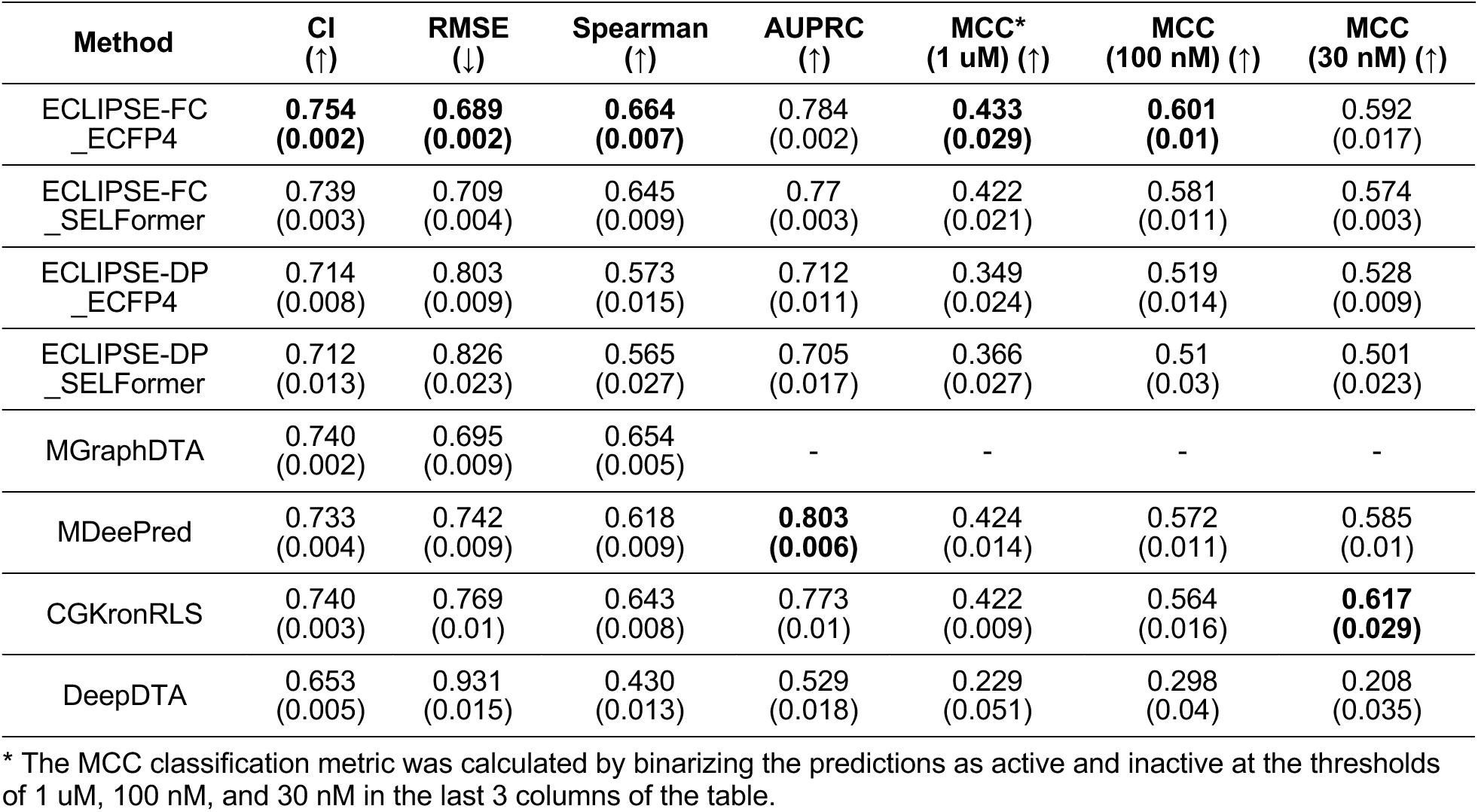
Performance comparison of ECLIPSE models with SOTA models on the filtered Davis kinase benchmark dataset. Standard deviations are given inside parentheses. The best performances are shown in bold font.

These outcomes are consistent with the random-split case of transferases family-based analysis, which was expected, given that the Davis kinase dataset is also randomly split. Table 1 and Figure 2 indicate that ECLIPSE-FC optimises binding-affinity prediction most effectively when train–test samples are similar, whereas dot-product variants achieve stronger out-of-sample generalisation on dissimilar splits. In ECLIPSE-DP, all trainable parameters reside in the HGT encoder, so interaction features must be captured within HGT; in ECLIPSE-FC, a learnable FC head before the output shifts part of the interaction modelling to the readout, allowing the encoder to prioritise node/edge-type relational structure.

### 2.4. Data type-centric ablation study

We performed an ablation study to investigate the effect of different graph node/edge types (i.e., biological components) on learning, using the three different data splits of the transferases dataset. For this analysis, we employed the ECLIPSE-DP_SELFormer model architecture. First, we trained the models on the KG encompassing all node and relationship types. Then, we systematically removed relationships associated with phenotypes, diseases, pathways, and DTIs. After each removal, a new model was trained and tested on the updated KG (using the exact same CPI data points). The models trained on the most basic KG only included protein-protein interactions (PPI) and compound-compound similarities (CCS). Each model was run five times, and the performances were averaged. Epsilon values were then calculated using the ASO test to assess the significance of performance differences between models (Supplementary Data Table 2). The performance results are provided in Figure 3 and Table S4.

**Figure 3.**
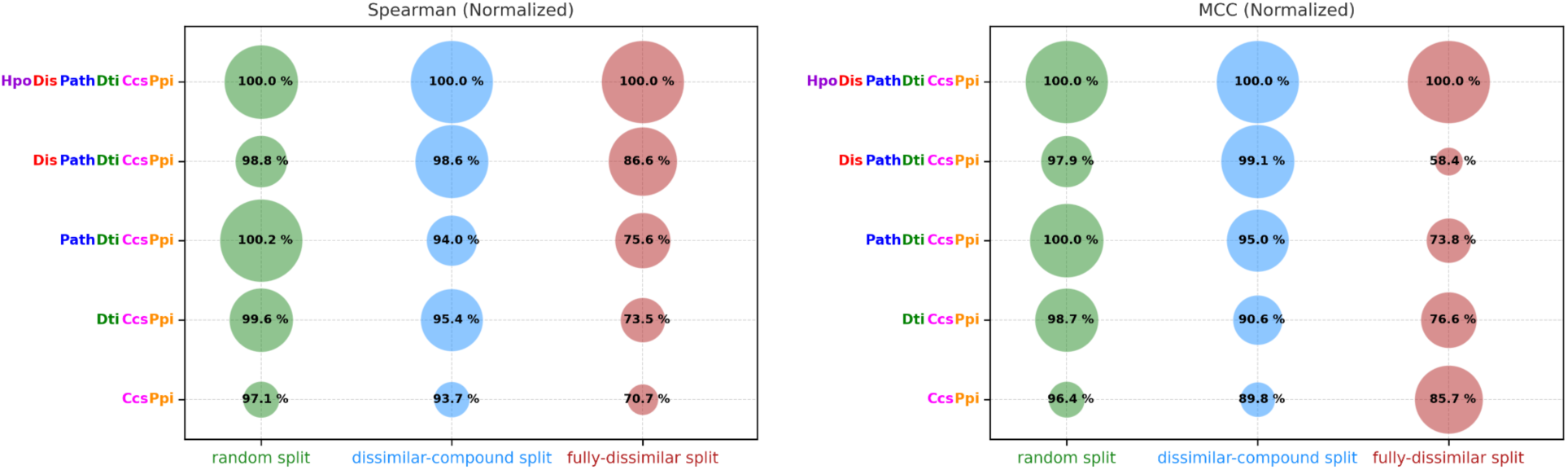
Dot plots of normalised test performance scores (Spearman and MCC) from the ablation study. Models differ from each other by input data types. The default model, i.e., the top row, incorporates all data modalities (Ppi: protein-protein interactions; Ccs: compound-compound similarities; Dti: drug-target interactions, including both biotech and small-molecule drugs; Path: protein-pathway associations from KEGG and Reactome; Dis: disease associations of proteins, drugs, and pathways from KEGG and EFO; Hpo: phenotype associations of proteins, pathways, and diseases from the Human Phenotype Ontology.

According to the Spearman and MCC values in Figure 3, omitting specific relationship types from the KG distinctly impacts ECLIPSE’s predictive performance across all splitting strategies (i.e., fully-dissimilar split “FDS”, dissimilar-compound split “DCS”, and random split “RS”).

Comparing the default ECLIPSE model (the top row in Figure 3) with the most basic model (the bottom row) shows a significant performance drop in all splits for both Spearman and MCC metrics (eps_min = 0.24 in FDS-Spearman and 0 in others; see Supplementary Data Table 2). For Spearman correlation, removing each relationship type gradually reduces performance, with the sharpest decline observed in the FDS scenario, especially when phenotype-and disease-related relationships are excluded. This highlights the relevance of disease-and phenotype-contextual information in predicting bioactivity for novel, distant compounds and targets. In the DCS and RS scenarios, the decline is comparatively milder, indicating complementary contributions of different relationships. MCC follows a similar pattern to Spearman for DCS and RS, yet diverges in the challenging FDS scenario—likely due to sensitivity when converting continuous predictions into binary (active/inactive) classification outcomes post hoc.

Overall, these results emphasise the value of incorporating multiple types of biological and biomedical relationships. By utilising rich structural and contextual information encoded in KGs, ECLIPSE demonstrates its ability to provide meaningful insights and accurate bioactivity predictions. These findings also suggest considerable potential for further improvement through incorporating additional biological relationships.

### 2.5. Exploring the Predictive Power of ECLIPSE in the Tails of the Bioactivity Distribution

Accurately predicting values at the lower and upper ends of the bioactivity range can be challenging for regression models, since these models typically focus on capturing overall trends in the data (Ribeiro & Moniz, 2020). However, reliable predictions in these regions, especially at the high end (e.g., pChEMBL value > 7), are critical for drug discovery. In this section, we examined how well ECLIPSE handles tails of the bioactivity distribution. We compared the predicted bioactivity distributions of ECLIPSE-DP_SELFormer and the baseline RF model with the actual distribution of test data points in the dissimilar-compound split of the transferases dataset. As illustrated in Figure 4a, the true distribution spans a broad pChEMBL range (4 to 11). The RF model predictions are concentrated around the median pChEMBL value (6.71), whereas ECLIPSE more closely recapitulates the empirical distribution.

**Figure 4.**
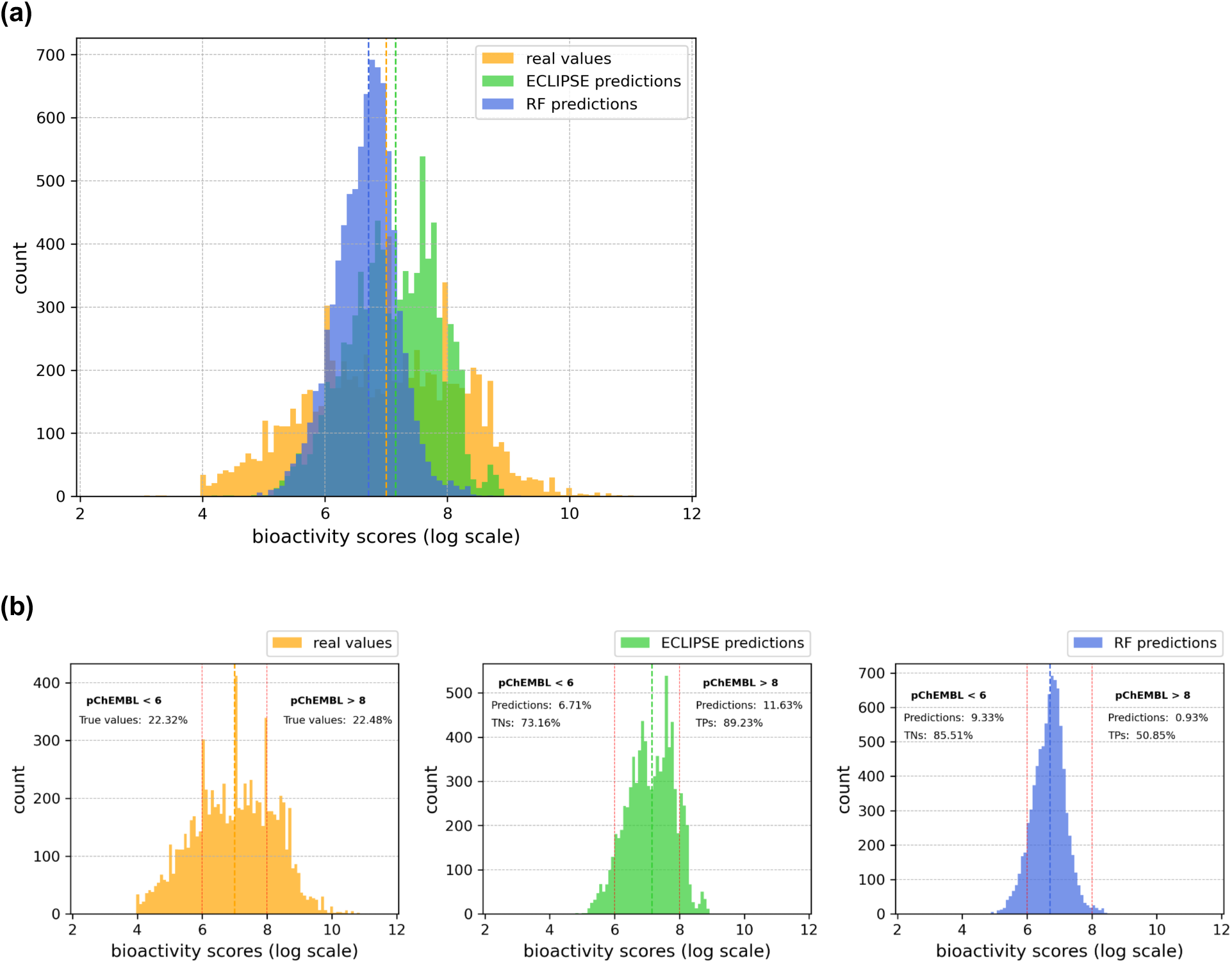
Bioactivity distribution histograms of real values, ECLIPSE predictions, and RF predictions, shown **(a)** collectively, and **(b)** separately, for test data points in the dissimilar-compound-split set of transferases family dataset. Bioactivity values are given in the log scale (pChEMBL values).

Figure 4b further presents quantitative comparisons in predicting values at the tails of the bioactivity distribution, defined as pChEMBL <6 and>8. In the true distribution (the left-most histogram in Figure 4b), each tail accounts for approximately 22% of the samples. Examining the distributions of predicted values from the ECLIPSE (the second histogram) and RF models (the third histogram), both models underrepresent these lower and upper ends. Specifically, RF captures 9.33% of samples with predicted pChEMBL <6 and only 0.93% with >8, showing a sharp drop-off in the high-activity range. ECLIPSE predicts 6.71% of values <6 and 11.63% >8, indicating a notably improved coverage in the high end. When evaluating model accuracy using an active/inactive threshold of pChEMBL: 6.97 (median of the training dataset), both models perform well in identifying true negatives (TNs) for pChEMBL <6 (ECLIPSE: 73%, RF: 86%).

However, ECLIPSE detects 89% of true positives (TPs) for pChEMBL >8, compared to RF’s 51%. Overall, while both models encounter challenges in accurately predicting values at the lower and upper ends of the bioactivity distribution, ECLIPSE displayed a better performance. This potential can be further enhanced by integrating data preprocessing strategies or adopting evaluation metrics tailored to imbalanced regression tasks ^54^.

### 2.6. Use-Case Study: Bioactivity Predictions for Druggable and Undruggable Proteins

To further evaluate the robustness and reliability of ECLIPSE, we conducted a use-case study centred on predicting compound interactions of both druggable and undruggable proteins. The aim is to assess whether the model distinguishes druggability profiles by learning meaningful patterns from compound–protein interactions and their contextual relationships, or simply memorises bioactivity values in relation to compound properties.

As a druggable protein, we selected PIM1 (Figure 5a). PIM1 is a proto-oncogene with serine/threonine kinase activity that plays a vital role in cell growth, survival, and apoptosis. It exerts its oncogenic effects through the regulation of MYC transcriptional activity, control of cell cycle progression, and inhibition of proapoptotic proteins via phosphorylation. Notably, abnormal elevation of PIM1 is associated with various types of cancer. It is a promising cancer drug target, particularly in prostate cancer ^55^. Figure 5a displays the PIM1 co-crystalized structure with a benzofuranone class PIM1 inhibitor (PDB ID: 5VUB). In our transferases training dataset (i.e., 138,297 data points in the dissimilar-compound-split), PIM1 has 3,019 data points with a mean bioactivity value of pChEMBL: 7.7.

**Figure 5.**
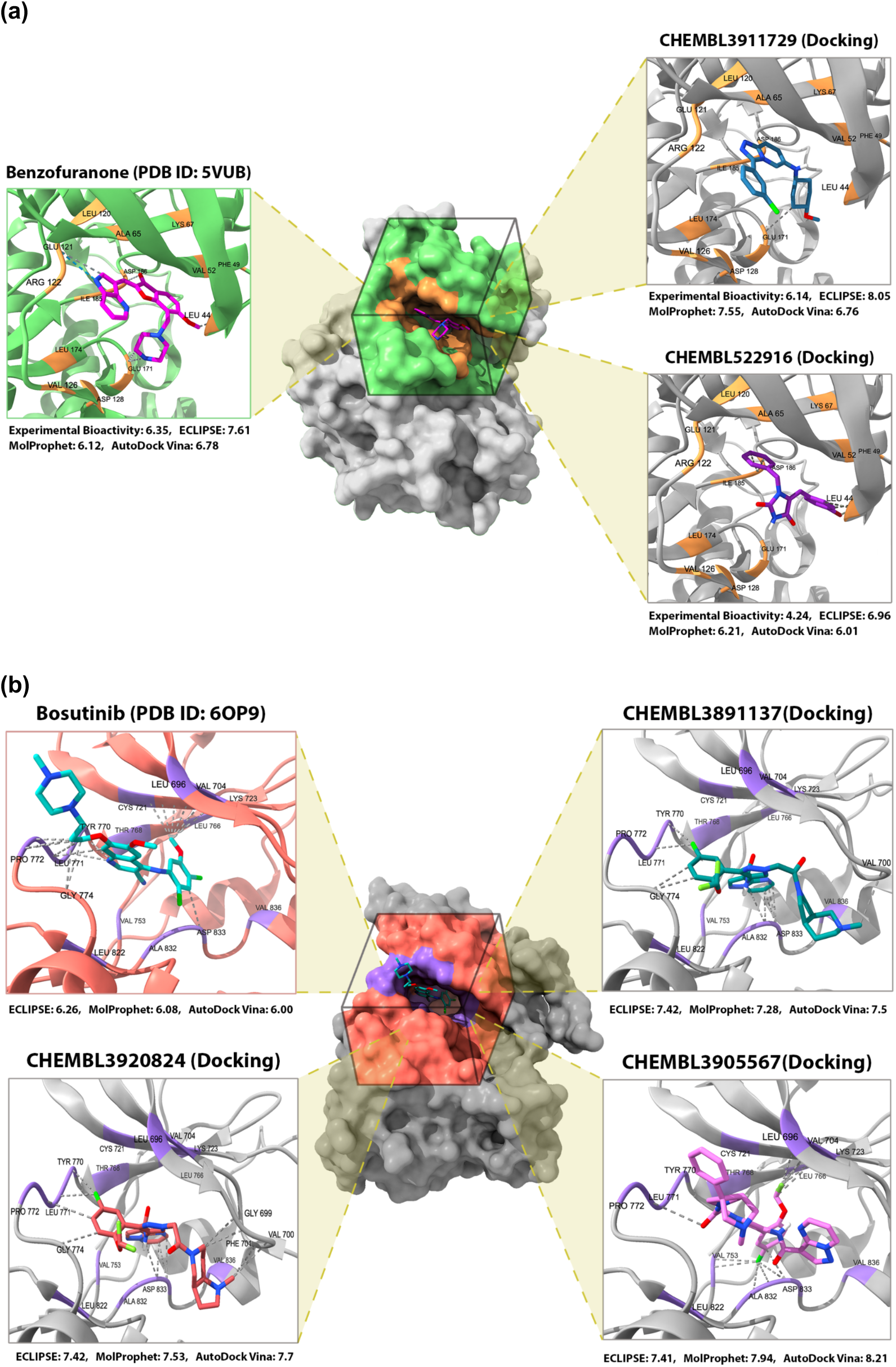
Co-crystalised 3-D structures of **(a)** PIM1 in complex with a benzofuranone, alongside best-scoring docking poses of CHEMBL3911729 and CHEMBL522916, and **(b)** HER3 in complex with bosutinib, along with best-scoring docking poses of CHEMBL3891137, CHEMBL3920824, and CHEMBL3905567. ECLIPSE predictions (pChEMBL values) are shown together with docking scores.

We selected HER3 as an undruggable protein (its protein structure is shown in Figure 5b; PDB ID: 6OP9). HER3, also known as ERBB3, is a pseudo-kinase member of the EGFR family, having a significant role in tumour progression and drug resistance. Unlike other kinases, its pseudo-kinase domain activates its partner HER family members, including HER2 and EGFR, through allosteric regulation. Despite its significance as a potential therapeutic target in various tumours, no HER3-directed therapies have received clinical approval to date ^56^. It has only 23 data points with a mean bioactivity value of pChEMBL: 5.9 in our training dataset, which is extremely low compared to PIM1.

#### 2.6.1. Druggability profiling of PIM1 - HER3 and structure-based assessment of HER3 hits

We assessed the bioactivity predictions of ECLIPSE-DP_SELFormer for PIM1 and HER3 targets against 422,617 compounds in our KG, using the model trained on the challenging dissimilar-compound split. To interpret the predictions, we binarised the continuous output values into active and inactive classes using the median pChEMBL value of the training set (i.e., 6.97) as the threshold, consistent with our prior classification protocol. Notably, 85% of PIM1 predictions were classified as active, whereas the rate for HER3 was 0.26% (1,083 compounds). These results align with the proportions observed in the training data: PIM1 was represented by 3,019 compound–target pairs, with 78% labelled as active, while HER3 had only 23 samples, with just 17% (4 out of 23) classified as active using the same threshold. Although the proportion of active compounds for PIM1 may seem inflated, it reflects a common characteristic of bioactivity datasets, such as ChEMBL, which are frequently enriched for active entries due to the design of screening libraries and publication bias ^57,58^. Moreover, PIM1, like many kinases, possesses a highly conserved ATP-binding pocket that can accommodate a wide range of small molecules, contributing to its binding promiscuity. It is a well-known property of kinases in general, making selective inhibition a major challenge in kinase drug discovery ^59^. The large proportion of predicted actives for PIM1 and the sparse hits for HER3 demonstrate the model’s ability to differentiate between druggable and undruggable targets.

While the extremely low proportion of predicted actives for HER3 reflects its status as an undruggable protein, the presence of a small set of positively predicted hits (1,083 compounds out of 422,617) raises interesting possibilities. Although some predictions may be false positives, others could represent previously unknown bioactive HER3 ligands. In the past, there have been similar instances, such as the KRAS protein, which was initially considered undruggable but later successfully targeted with specific types of compounds ^60^.

To further evaluate the matter, we analysed the three most active interacting compound predictions by ECLIPSE for HER3, by conducting docking analyses using two complementary methods: the AI-based docking tool of the MolProphet platform and the classical physics-based docking engine AutoDock Vina. Figure 5b and Table S5 present the predicted pChEMBL values from the ECLIPSE model, estimated binding affinities from MolProphet, and dissociation constant (pKd) values derived from AutoDock Vina by converting the binding free energies of the best-scoring poses to a comparable bioactivity scale. For comparison, data for Bosutinib, the reference molecule from the co-crystallised PDB structure of HER3, is also provided. The analysis revealed that the top three hit molecules (CHEMBL3891137, CHEMBL3920824, and CHEMBL3905567) yielded consistent scores across all three methods, demonstrating stronger predicted activity than the reference molecule Bosutinib, which had bioactivity prediction and docking scores around 6 (corresponding to 1 µM). Figure 5b illustrates the best-scoring docking poses of these three compounds alongside Bosutinib within the ATP-binding pocket of the HER3 kinase domain (PFAM: PF07714), as determined by AutoDock Vina. Each compound forms multiple close contacts with surrounding residues (van der Waals overlap ≥ –0.4 Å) and exhibits greater binding site occupancy and shape complementarity than Bosutinib. Their improved fit within the ATP-binding cleft, coupled with consistently high scores across both AI-based and physics-based docking methods, supports their potential as viable HER3 ligands.

These findings warrant further experimental investigation to confirm their activity and therapeutic potential.

#### 2.6.2. Compound-centric analysis of PIM1 predictions

In this subsection, our goal is to assess model consistency across training and test sample predictions and to investigate whether certain compound groups or structures influence the prediction capability of ECLIPSE. We focused on the PIM1 protein for this analysis.

First, we evaluated the performances of the ECLIPSE-DP_SELFormer and the baseline RF model for predictions on the training and test set compounds of the PIM1 protein. The models yielded the following Spearman’s correlation scores: 0.75 (ECLIPSE) and 0.99 (RF) for training (3,019 samples); 0.74 (ECLIPSE) and 0.47 (RF) for test (36 samples). The nearly perfect score for the RF model on the training samples, followed by a significant drop in performance on the test samples, indicates memorisation and overfitting to the training data. In contrast, the highly consistent scores for the ECLIPSE model across both training and test data underscore learning as desired.

Secondly, we investigated the compound diversity of the PIM1 bioactivity dataset to evaluate the model’s capacity for correctly predicting a diverse range of compound structures versus being limited to a narrow set of easily predictable structures. To achieve this, we clustered the 3,019 training compounds based on their pairwise Tanimoto similarities using the Butina clustering function in RDKit, with a cut-off value 0.5. Figure 6 displays the prediction error box plots of clusters with more than five compound members. To obtain prediction errors, we computed the differences between the predicted and actual bioactivities of each sample, expressed as pChEMBL units. In this plot, blue bars represent compound clusters with overestimated bioactivity values, while red bars correspond to clusters with underestimations. Upon analysing the median prediction errors across these compound clusters, we observed that a significant portion of the clusters (63 out of 83) fall within the prediction error range of-1 to 1. This indicates the effectiveness of our ECLIPSE model in accurately predicting bioactivity across diverse compound groups, rather than being limited to a narrow subset.

**Figure 6.**
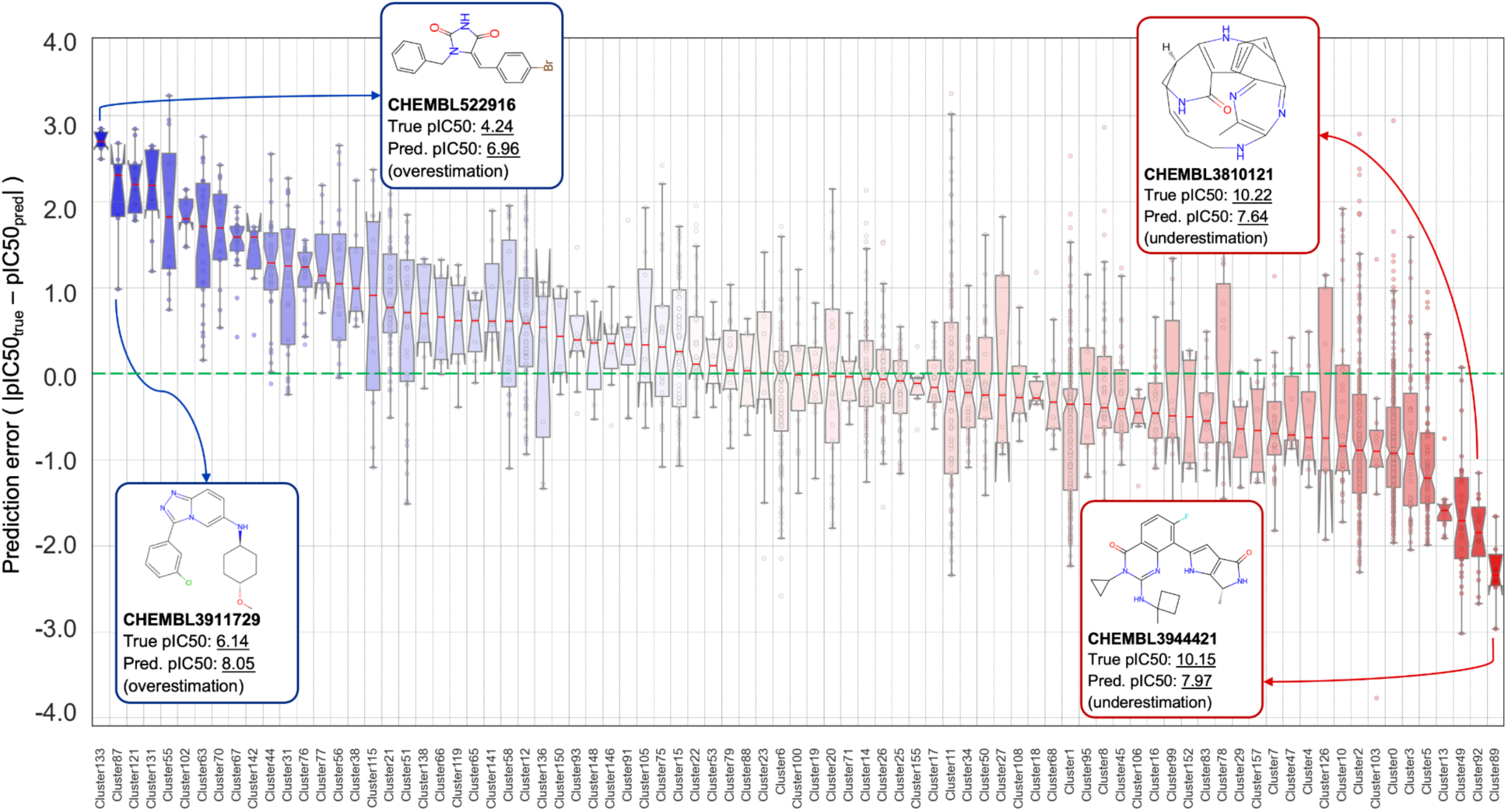
Bioactivity prediction error box plots of compound clusters for the PIM1 protein in the dissimilar-compound-split of the transferases training dataset. The plot also displays four representative compounds from the dataset, belonging to over-and under-estimated clusters, along with their experimental (“true”) and predicted (“pred.”) bioactivities (i.e., pIC50).

While evaluating the median prediction errors beyond the acceptable range (i.e., <-1 and> 1), we observe a higher number of overestimations (15 clusters) compared to underestimations (5 clusters). To closely examine these clusters, we picked representative compounds from clusters 87 (CHEMBL3911729) and 133 (CHEMBL522916) for overestimation, and from clusters 89 (CHEMBL3944421) and 92 (CHEMBL3810121) for underestimation. Figure 6 displays these representative compounds alongside their cluster information and experimental and predicted bioactivities, exemplifying the molecular characteristics of the respective clusters.

To assess the structural plausibility of ECLIPSE’s overestimated predictions for PIM1, we performed molecular docking on CHEMBL3911729 and CHEMBL522916 (Figure 5a).

MolProphet estimated bioactivities of 7.55 and 6.21, and AutoDock Vina yielded pKd values of ∼6.76 (–9.21 kcal/mol) and ∼6.01 (–8.19 kcal/mol), respectively, for these compounds. Although both docking methods scored lower values than ECLIPSE’s bioactivity predictions (8.05 and 6.96, respectively), their estimates remained notably higher than the experimentally reported IC₅₀-based pChEMBL values (6.14 and 4.24), suggesting stronger binding potential than observed in vitro. Both molecules contain halogen substituents (e.g., Br, Cl, and F), a feature commonly exploited in kinase inhibitor design to improve potency through enhanced hydrophobic contacts and halogen bonding ^61^. Given the prevalence of halogen-containing groups in numerous bioactive compounds, our model may have learned to associate these groups with increased bioactivity, resulting in higher predicted values. As displayed in Figure 5a, a similar-though less pronounced-overestimation was observed for benzofuranone, the co-crystallized ligand of PIM1 (PDB ID: 5VUB). Its planar heterocyclic scaffold, a common feature in ATP-competitive kinase inhibitors ^62^, may contribute to the elevated predicted activity, despite the absence of halogens. It is also important to note that the experimental values of these compounds are derived from IC₅₀ assays, which are influenced by assay-specific conditions such as ATP concentration. For ATP-competitive inhibitors, high ATP levels can substantially increase IC₅₀ (corresponding to lower experimentally measured pChEMBL values) relative to true binding affinity, potentially contributing to the observed discrepancies. While we acknowledge that both docking and ECLIPSE remain in silico approaches and cannot independently verify true bioactivity, the consistency across methods and the structural compatibility of the ligand poses suggest that these compounds could merit experimental re-evaluation.

For the compounds with underestimated bioactivity (Figure 6), the predicted values were high but still lower than the experimentally observed ones. These molecules feature complex structures with unique substituents and display exceptionally strong bioactivity, which is both uncommon in the training dataset and located at the extreme end of the distribution. This rarity limits the model’s ability to learn their distinctive features and accurately capture bioactivity.

Taken together, our findings show that ECLIPSE successfully captures structure–activity relationships across a broad spectrum of chemically diverse scaffolds, an essential quality for real-world virtual screening applications. While a small subset of clusters exhibited systematic over-or under-estimations, these appear to be isolated cases, likely influenced by subtle chemical characteristics or dataset-specific variations.

## 3. Conclusion

This study introduces ECLIPSE, a novel heterogeneous graph representation learning-based framework for providing systems-level representations of proteins and compounds and subsequent prediction of their interactions. Utilising large-scale biomedical knowledge graphs constructed by the CROssBAR system, ECLIPSE effectively captures the complex relationships between genes/proteins, pathways, diseases, phenotypes, drugs, and compounds. The heterogeneous graph transformer architecture is employed to learn from these diverse biomedical relationships and generate integrative representations that carry rich structural and contextual information encoded in KGs. Our benchmarking experiments on the large-scale ProtBENCH transferases dataset demonstrate the superiority of ECLIPSE over baseline models, showcasing its adaptability and potential for realistic CPI prediction scenarios. It also highlights the generalizability of ECLIPSE with improved predictions for previously unseen data, addressing one of the major bottlenecks in drug discovery. Furthermore, the outstanding performance of ECLIPSE against state-of-the-art models on the Davis kinase bioactivity benchmark dataset validates its effectiveness as a KG-based learning approach.

At this point, it is crucial to emphasize that the Davis kinase benchmark is a medium-scale dataset with significantly fewer datapoints compared to ProtBENCH family-specific datasets (i.e., 9,085 datapoints in Davis, as opposed to 146,677 datapoints in the transferases family subset of ProtBENCH). Furthermore, the random splitting of the dataset simplifies the prediction task, potentially resulting in over-optimistic results. As a result, it lacks representativeness and may not align well with real-world scenarios in CPI prediction modeling, despite its common usage. Hence, we strongly advocate for the use of ProtBENCH large-scale family datasets (either the graph-integrated filtered version used here or the original version in the GitHub repository of our previous study, available at https://github.com/HUBioDataLab/ProtBENCH), which consider multiple difficulty levels of the task by applying alternative data splitting strategies.

In the ablation study, we explored the impact of various graph node/edge types on learning and found that ECLIPSE effectively leverages structural and contextual information encoded in KGs. The model performance is notably improved with the addition of new data layers to the graph, particularly in challenging cases (i.e., on the fully-dissimilar-split dataset).

While the ECLIPSE system exhibits limitations in precisely predicting values at the tails of the bioactivity distribution, a known issue with regression-based CPI prediction models, it demonstrates proficiency in correctly classifying these values into active or inactive classes. The outcomes of the use-case study on the predictions of druggable (PIM1) and undruggable proteins (HER3) further support its reliability.

To enable reproducibility and community reuse, we have released the ECLIPSE codebase, datasets, trained models, and benchmark results in our public GitHub repository at https://github.com/HUBioDataLab/ECLIPSE. This codebase can be used to train models for different target protein families and subsequently employed for novel ligand-target interaction predictions.

There are several promising avenues for further exploration and development. Firstly, expanding the application of ECLIPSE to additional protein superfamilies is valuable, as it can enhance the model’s versatility and effectiveness across a broader spectrum of biological contexts. Secondly, the integration of additional data sources into the KGs could enhance the predictive power of ECLIPSE. In this context, integrating new types of nodes and edges to the KG, such as cell-based information including drug sensitivity measurements, gene expression profiles, mutations, together with biological ontologies, side effects and toxicity profiles of drugs and drug candidate compounds, and metabolomics data could provide a richer biological and chemical representation. Furthermore, the incorporation of an iterative active learning process during model training could facilitate more efficient and targeted data acquisition, leading to better model performance, especially for underrepresented targets or protein families, via yielding data augmentation. One crucial aspect to focus on is the interpretability of the proposed method to unravel the underlying mechanisms and features contributing to the decision-making process.

In summary, ECLIPSE integrates multi-layer biomedical knowledge with learned embeddings in a knowledge-graph–based representation-and-prediction framework. Together with the planned follow-up studies, this systems-level approach can systematically prioritise mechanism-grounded candidates across indications and support the discovery of effective therapies.

## 4. Methods

### 4.1. Dataset Construction

To develop robust and effective GNN-based CPI prediction models, the choice of input graph data is as crucial as the selection of the algorithm and input features. In this study, we utilised CROssBAR knowledge graphs ^18^, comprising genes/proteins, drugs/compounds, disease/phenotype terms and pathways/biological processes represented as nodes, with relationships between them annotated as edges. The following step-by-step operation was executed to construct our heterogeneous biological KG data to be used in the training and testing of the ECLIPSE system:

1) We performed a bulk search via the CROssBAR web service (https://crossbar.kansil.org/) to obtain individual KGs for all reviewed human proteins in the UniProt database (i.e., 20,173 protein entries in UniProtKB/Swiss-Prot) ^39^, by querying each protein entry with parameters; number of experimentally validated drugs targeting the protein (num_of_drugs) = 100, number of experimentally validated compounds targeting the protein (num_of_compounds) = 100, and number of predicted drugs/compounds (predictions) = 0, where other parameters were set as default.
2) These CROssBAR KGs, which include query proteins, other proteins/drugs/compounds interacting with those query proteins, their associated pathways, diseases, and phenotypes obtained in Step 1 were merged to create an extensive KG encompassing all human proteins and their associated relationships. Duplicate nodes and edges were removed.
3) For the training and testing of the models, we used protein family-specific ProtBENCH bioactivity datasets ^50^, i.e., bioactivity measurements of compound-target protein interactions based on pChEMBL values. They were filtered so that only bioactivity data points belonging to human proteins in our KG (Step 2) were included. Compounds that have bioactivity data with these proteins (but not included in the KG) were manually integrated as additional compound nodes.
4) We excluded all compound-protein edges from the KG to prevent data leakage and limited their usage solely as training and test labels, meaning that these edges are not used during the message passing procedure during graph learning. It’s also worth noting that we maintain drug and compound nodes as distinct node types, and drug-protein interactions are not considered in our training/testing data. Hence, while we omitted compound-protein edges, drug-protein edges were retained in the KG.
5) We incorporated compound-compound similarity edges. Pairwise compound similarities were computed using the simsearch function of the Chemfp Python package ^63^. Pairwise similarities with scores higher than 0.5 were added to the graph as edges between compound nodes. The aim behind this step was to maintain graph connectivity following the exclusion of protein-compound edges (i.e., reconnecting the compound nodes to the graph).
6) To construct regression-based CPI prediction models optimized specifically for the transferases protein family, we labeled the train and the test bioactivity edges (i.e., compound-target protein) on the KG. This process was carried out for three different train/test data split versions, i.e., fully-dissimilar-split (FDS), dissimilar-compound-split (DCS), random-split (RS), of the ProtBENCH transferases dataset. Train/test data point counts are 139,292/7,385 for FDS; 138,297/8,380 for DCS; and 137,727/8,950 for RS.

We used the filtered Davis kinase benchmark dataset, following the same configuration as described in the MDeePred study ^15^, to compare ECLIPSE with the state-of-the-art (SOTA) models in the literature. We integrated the filtered Davis dataset into our KG using the same process applied for the ProtBENCH transferases bioactivity dataset (described above). This KG consisted of 7,567 training and validation, and 1,518 test data points, which was 7,600 and 1,525 in the original MDeePred study (9,125 data points in total, with the train/validation/test ratio of 4:1:1). The difference was due to the absence of KGs for three proteins (UniProt IDs: Q07785, P62344, and P9WI81) in CROssBAR since they belong to Plasmodium falciparum and Mycobacterium tuberculosis organisms.

Table S6 presents the statistics of our integrated CROssBAR KG, serving as the basis for mapping training/test samples in both the transferases and the Davis datasets.

### 4.2. The Representation (Features) of Graph Nodes

Graph representation learning algorithms enable knowledge transfer between nodes by employing their respective feature vectors. Although it is possible to initialise these feature vectors randomly or to use trivial solutions such as one-hot encodings of pre-determined features, utilising learned representations (embeddings) to capture hidden patterns of nodes may increase the success rate of prediction models. Below, we explained the representation approach employed for each node type to construct the input node embeddings:

-*Proteins* were represented by concatenated vectors of transformer-avg embeddings (vector size: 768), apaac descriptors (vector size: 80), and k-sep_pssm descriptors (vector size: 400). For the construction of transformer-avg embeddings, we used BERT algorithm-based model developed by Rao et al. ^64^. Apaac descriptors (i.e., “Amphiphilic Pseudo Amino Acid Composition”—physicochemical properties of amino acids) were generated using iFeature stand-alone tool ^65^, and k-sep_pssm descriptors (i.e., “k-separated-bigrams position specific scoring matrix”—encoding evolutionary relationships between proteins) were obtained via the POSSUM web server ^66^.
-*Compounds/Small molecule drugs* were encoded by SELFormer embeddings (vector size: 768). SELFormer ^49^ is a transformer-based NLP model that employs large-scale pre-training on 2 million molecules using their SELFIES notations ^67^, enabling the acquisition of flexible and high-quality molecular representations. Additionally, in our alternative models, we used ECFP4 fingerprints ^52^ with a size of 1024 to represent compound and drug nodes.
-*Biotech drugs* were represented by protein embeddings (i.e., transformer-avg embeddings), as they are composed of amino acid sequences.
-*KEGG pathway* representations (vector size: 200) were generated using the TransE embedding method via the BioKEEN library ^68^, which utilizes a gene-pathway association network to generate representations. For *Reactome pathways*, we used pre-calculated Bioteque KG embeddings (vector size: 128) ^19^ generated on gene-pathway-disease metapath using a random walk method.
-*HPO phenotype term* embeddings (vector size: 160) were retrieved from the CADA tool^69^. CADA is a node2vec-based embedding method using a gene-phenotype association network as the input graph that includes disease-level annotations and clinical cases-level annotations.
-*Disease* embeddings (vector size: 100) were obtained using the doc2vec method based on the PrimeKG ^70^ disease definitions.

### 4.3. Model Design and Architecture

We adopted the heterogeneous graph transformer (HGT) architecture ^47^ at the core of our ECLIPSE framework since this algorithm suits the problem of learning meaningful representations of nodes in a highly heterogeneous graph, based on their intrinsic properties (feature vectors) and relationships.

As shown in Figure 1, the HGT module within the ECLIPSE system incorporates information from source nodes in the CROssBAR KG with the input features described in Methods Section 4.2. It then generates contextualized representations for target nodes through a learning process involving heterogeneous mutual attention, heterogeneous message passing, and target-specific aggregation steps, as outlined below:

<u>1) Heterogeneous mutual attention:</u> HGT introduces a new mechanism for attention calculation that considers the meta-relations between nodes, which is inspired by the Transformer architecture ^71^. Given a graph with the input node features 𝐻^(𝑙−1)^, each node type’s features are first projected into a common hidden space using a type-specific linear transformation followed by a ReLU activation to ensure uniform dimensionality. The attention mechanism then computes a Key vector *K* for the source node *s* (Equation 1) and a Query vector *Q* for the target node *t* (Equation 2) through separate linear projections, forming the basis for relation-aware attention calculation.

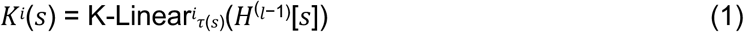

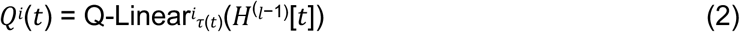

Then, instead of directly taking the dot product of the Query and Key vectors like the vanilla Transformer, it uses different weight matrices (𝑊^𝐴𝑇𝑇^) for each edge type (𝜙(𝑒)) to calculate the attention matrix for *h* heads, allowing it to capture different semantic relationships between nodes. Additionally, a prior tensor (𝜇) is introduced to represent the general significance of each meta-relation triplet (<𝜏(𝑠),𝜙(𝑒),𝜏(𝑡)>) since not all the relationships contribute equally to the target nodes. The prior tensor is divided by the square root of the vector dimension per head (√ 𝑑), which serves as an adaptive scaling factor for attention (Equation 3).

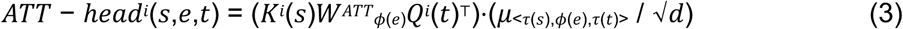

To obtain the attention vector of each node pair, multiple attention heads are concatenated.

Then, for each target node *t*, the attention vectors are gathered from its neighboring source nodes *N(t)*, and the softmax function is applied to ensure attention weights sum up to 1 across the source nodes (Equation 4).

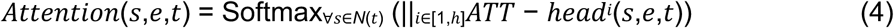

<u>2) Heterogeneous message passing:</u> Following the calculation of mutual attention (Equation 4), HGT performs message passing from source nodes to target nodes. Similar to attention, this process incorporates the meta relations of edges into the message passing to address the distribution differences of different node and edge types. For each pair of nodes *e = (s,t)*, HGT calculates a multi-head message by projecting the source node *s* into message vectors M and using a matrix (𝑊^𝑀𝑆𝐺^) to incorporate edge dependency (Equation 5).

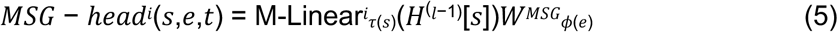

Multiple message heads (*h*) are concatenated to obtain the message for each node pair (Equation 6).

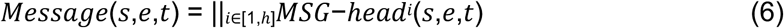

<u>3) Target-specific aggregation:</u> After the calculation of heterogeneous mutual attention (Equation 4) and message passing (Equation 6), HGT aggregates the information from the source nodes to the target node. The attention vectors obtained from the softmax procedure in the first step (𝐴𝑡𝑡𝑒𝑛𝑡𝑖𝑜𝑛(𝑠,𝑒,𝑡)) serve as weights to average the corresponding messages from source nodes (𝑀𝑒𝑠𝑠𝑎𝑔𝑒(𝑠,𝑒,𝑡)) to get the updated vector 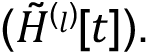 This aggregation process is performed for all target nodes, incorporating information from their neighboring source nodes of different feature distributions (Equation 7).

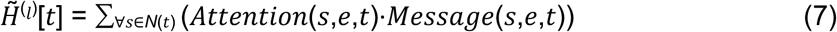

The updated vector for each target node is then mapped back to its type-specific distribution using a linear projection followed by a non-linear activation and residual connection (Equation 8). The output vector for the target node *t* is computed as follows:

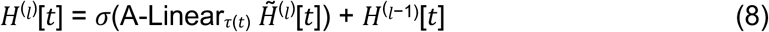

By stacking multiple HGT blocks for *L* number of layers, the model enables each node to reach a large proportion of nodes with different types and relationships in the full graph.

Once the embeddings are learned via the HGT module, the ECLIPSE system performs an edge-level regression task to predict CPIs. In the default setup, it computes the dot product of the linear-transformed learned embeddings of compound and proteins for each pair in the training set, producing predicted bioactivity values (i.e., pChEMBL scores). In addition, an alternative model variant, referred to as ECLIPSE-FC, concatenates the linearized compound and protein embeddings and processes them through a deep neural network, a multi-layered perceptron. This network consists of multiple fully-connected (FC) layers, where each layer applies a linear transformation, followed by batch normalization and a ReLU activation, before passing the output to the next hidden layer. The final output layer produces CPI predictions (i.e., pChEMBL scores) through a final linear projection. During the training of the models, the loss is calculated based on the true and predicted bioactivity values of the compound-protein pairs using the mean squared error (MSE) loss function (i.e., L2 loss). Subsequently, the weights of the HGT layers are updated through backpropagation, using the Adam optimizer. This iterative process continues until the loss converges. Following the training phase, the models are utilised to generate CPI predictions for the samples in the validation and test sets. These predictions are then used to evaluate the performance of the models.

### 4.4. Performance Evaluation

To assess the performance of models trained and tested on the ProtBENCH transferases family dataset, we employed Spearman rank correlation, root mean square error (RMSE), and Matthews correlation coefficient (MCC) as scoring metrics. Formulae of the corresponding metrics are given below:

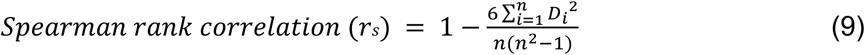

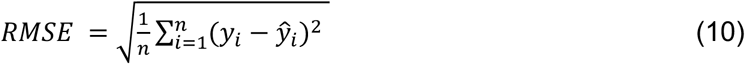

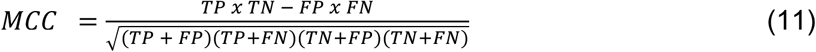

where 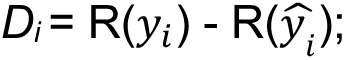 denotes the difference between ranks of true (𝑦_i_) and predicted 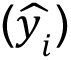 values of samples with the dataset size *n*. TP, TN, FP, and FN represent the total counts of true positive, true negative, false positive, and false negative predictions, respectively.

To compute TP, TN, FP, and FN for the calculation of MCC, we binarized the continuous prediction scores into active (1) and inactive (0) classes. The threshold for classification was set to the median pChEMBL value within each training dataset: 6.975 for RS, 6.97 for DCS, and 6.99 for FDS. Compound–target pairs with bioactivity values above the median were labeled as active, while those equal to or below the median were labeled as inactive.

The models trained and tested on the filtered Davis kinase benchmark dataset were evaluated via 5-fold cross validation (CV) using the same metrics in the MDeePred study ^15^, which include RMSE, Spearman, MCC, average area under the precision-recall curve (AUPRC), and concordance index (CI) scores. For the MCC score, three different threshold values (i.e., 1 uM, 100 nM, and 30 nM) were used to binarize the regression-based CPI predictions. For computing the average AUPRC score, ten interaction threshold values from the pKd interval [6 M, 8 M] were considered to binarize pKds into true class labels. CI for a set of paired data (i.e., compound-protein pairs) represents the probability that the predictions for two randomly selected pairs are correctly ordered based on their respective labels (i.e., pKd values). It ranges from 0 to 1, where 1 indicates perfect agreement between the predicted and observed rankings^72^. The formula used for the calculation of CI is provided below:

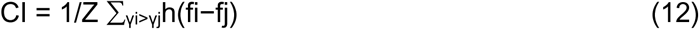

where fi and fj represent the predicted binding affinity values for the larger affinity γi and the smaller affinity γj, respectively. Z is a normalisation constant equal to the number of pairs, and h(u) is the Heaviside step function ^73^ that returns 1.0 for u > 0, 0.5 for u = 0, and 0.0 for u < 0.

### 4.5. Molecular Docking Experiments

For the docking analyses of PIM1 and HER3 proteins with their respective ligands (Bosutinib - CHEMBL288441-, CHEMBL3891137, CHEMBL3920824, and CHEMBL3905567 for HER3; Benzofuranone-CHEMBL2147770-, CHEMBL3911729, and CHEMBL522916 for PIM1), we utilized their crystallized structures from the Protein Data Bank (Berman et al., 2000) (PDB IDs: 5VUB for PIM1 and 6OP9 for HER3, see Figures 5a and 5b). We employed two docking methods: the traditional AutoDock Vina approach and the AI-based docking model of the MolProphet platform.

For AutoDock Vina, we used the SwissDock web service (https://www.swissdock.ch/). We provided the SMILES representations of the compounds retrieved from the ChEMBL database ^74^ and the corresponding PDB structures of the proteins. All heteroatoms were removed from the protein structures before docking. The search box was defined based on the mean coordinates of interacting atoms in the crystallized structures, as described in the study by Rifaioglu et al. ^75^. The mean coordinates are as follows: x: 40.57, y: 2.38, z: 1.34 for PIM1 (co-crystallized with ligand 8GU, PDB ID: 5VUB) and x:-50.53, y:-13.65, z: 12.93 for HER3 (co-crystallized with ligand DB8, PDB ID: 6OP9). The search box size was set to 30×30×30 Å, and the sampling exhaustivity was set to the default value of 4. To convert binding free energies (ΔG) estimated by AutoDock Vina into dissociation constant values (pKd), we utilized the NovoPro online conversion tool (https://www.novoprolabs.com/tools/deltag2kd). The tool applies the formula:

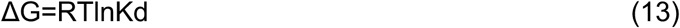

where ΔG is Gibbs free energy (kcal/mol), R is the universal gas constant (1.987 cal/(mol·K)), T is the temperature (298 K under standard conditions), and Kd is the dissociation constant (mol/L).

For MolProphet, we used its AI Docking function via its web platform (https://app.molprophet.com/). This module integrates geometric deep learning to learn structural features of both protein binding pockets and small molecules, and employs reinforcement learning to sample flexible receptor conformations while optimizing ligand binding poses to predict the minimum binding free energy. This AI-driven approach enables target-aware predictions, making it particularly well-suited for high-throughput virtual screening ^76^. We provided PDB IDs for target proteins and SMILES strings for ligands; the tool autonomously identified likely binding pockets and predicted interactions accordingly. Docking results and binding interactions were visualized using UCSF ChimeraX (version 1.10) (see Figure 5a and 5b).

## 5. Data and code availability

The source code, datasets, and the results of this study are available at https://github.com/HUBioDataLab/ECLIPSE.

## 6. Author Contributions

TD conceptualised the study and designed the general methodology. HAG prepared the datasets and the codebase, designed, implemented, trained, tuned and evaluated the ECLIPSE models, and conducted docking analysis. HAG and TD prepared the figures and wrote the manuscript. HAG prepared the code repository. TD supervised the overall study. Both authors approved the manuscript.

## 7. Conflicts of interest

There are no conflicts to declare.

## 8. Acknowledgements

The authors thank Dr. Taylan Güvenilir from the Faculty of Fine Arts at Gaziantep University, Turkey, for his valuable contributions and assistance in creating the workflow illustration and integrated molecular docking visualisations featured in this study.

## Notes

### Competing Interest Statement

The authors have declared no competing interest.

