## Supplementary Information for "ECLIPSE: Exploration of Complex Ligand-Protein Interactions through Learning from Systems-level Heterogeneous Biomedical Knowledge Graphs"

**Table S1.** Model optimisation for the given hyperparameters with their respective ranges.

| **Hyperparameter Name** | **Selected Value(s)** |
|---|---|
| hidden channels | 16, 32, 64, 128, 256 |
| learning rate | 0.00001, 0.00005, 0.0001, 0.0005, 0.001, 0.005 |
| head number | 4, 8, 16, 32, 64 |
| layer number | 1, 2, 3 |
| train batch size | 128, 256, 512, 1024 |
| epoch number* | 5 |
| weight decay | 0, 0.0001, 0.0005, 0.001, 0.005 |
| dropout ratio (only for FC models) | 0.1, 0.3, 0.5 |
| FC layer number (only for FC models) | 2, 3, 4 |

*Following the determination of these hyperparameters, we performed additional fine-tuning for the epoch number.

**Table S2.** Hyperparameters of the finalised ECLIPSE models on random-split (RS), dissimilar-compound-split (DCS), and fully-dissimilar-split (FDS) datasets of transferases.

| Dataset Split | Model Name | Epoch Number | Hidden Channels | Learning Rate | Head Number | Layer Number | Batch Size | Weight Decay | Dropout |
|---|---|---|---|---|---|---|---|---|---|
| RS | ECLIPSE-DP_ SELFormer | 250 | 128 | 0.001 | 16 | 1 | 1024 | 0 | - |
| RS | ECLIPSE-DP_ ECFP4 | 50 | 128 | 0.001 | 16 | 1 | 1024 | 0 | - |
| RS | ECLIPSE-3FC_SELFormer | 300 | 128 | 0.001 | 32 | 1 | 1024 | 0 | 0.1 |
| RS | ECLIPSE-3FC_ECFP4 | 50 | 128 | 0.001 | 32 | 1 | 1024 | 0 | 0.3 |
| RS | DP_SELFormer | 300 | 256 | 0.001 | - | - | 1024 | 0 | - |
| RS | DP_ECFP4 | 80 | 256 | 0.001 | - | - | 1024 | 0 | - |
| RS | 3FC_SELFormer | 300 | 256 | 0.001 | - | - | 1024 | 0 | 0.1 |
| RS | 3FC_ECFP4 | 150 | 256 | 0.001 | - | - | 1024 | 0 | 0.1 |
| DCS | ECLIPSE-DP_ SELFormer | 25 | 32 | 0.0001 | 8 | 3 | 512 | 0 | - |
| DCS | ECLIPSE-DP_ ECFP4 | 15 | 32 | 0.0001 | 8 | 3 | 256 | 0 | - |
| DCS | ECLIPSE-2FC_SELFormer | 20 | 32 | 0.0001 | 8 | 3 | 512 | 0 | 0.1 |
| DCS | ECLIPSE-2FC_ECFP4 | 20 | 32 | 0.00005 | 8 | 2 | 256 | 0.005 | 0.3 |
| DCS | DP_SELFormer | 50 | 256 | 0.00001 | - | - | 1024 | 0.005 | - |
| DCS | DP_ECFP4 | 30 | 256 | 0.00001 | - | - | 1024 | 0.005 | - |
| DCS | 2FC_SELFormer | 30 | 256 | 0.00001 | - | - | 1024 | 0.005 | 0.1 |
| DCS | 2FC_ECFP4 | 30 | 256 | 0.00001 | - | - | 1024 | 0.005 | 0.1 |
| FDS | ECLIPSE-DP_ SELFormer | 20 | 32 | 0.0001 | 8 | 2 | 512 | 0 | - |
| FDS | ECLIPSE-DP_ ECFP4 | 20 | 32 | 0.0001 | 8 | 2 | 512 | 0 | - |
| FDS | ECLIPSE-2FC_SELFormer | 10 | 64 | 0.0001 | 8 | 2 | 1024 | 0.0001 | 0.3 |
| FDS | ECLIPSE-2FC_ECFP4 | 10 | 64 | 0.0001 | 8 | 2 | 1024 | 0.005 | 0.1 |
| FDS | DP_SELFormer | 20 | 32 | 0.00001 | - | - | 256 | 0 | - |
| FDS | DP_ECFP4 | 20 | 128 | 0.00001 | - | - | 256 | 0 | - |
| FDS | 2FC_SELFormer | 40 | 64 | 0.0001 | - | - | 1024 | 0.005 | 0.3 |
| FDS | 2FC_ECFP4 | 20 | 64 | 0.0001 | - | - | 1024 | 0.005 | 0.3 |

**Table S3.** Performance score comparison of ECLIPSE and baseline models on the fully-dissimilar, dissimilar-compound, and random splits of the transferases test bioactivity dataset. The best performance for each split is shown in bold font.

| Method | Data Split | RMSE | Spearman | MCC |
|---|---|---|---|---|
| RF_SELFormer | RS* | 0.828 | 0.765 | 0.576 |
| RF_ECFP4 | RS | **0.643** | **0.861** | **0.695** |
| DP_SELFormer | RS | 0.929 | 0.687 | 0.492 |
| DP_ECFP4 | RS | 0.709 | 0.837 | 0.666 |
| 3FC_SELFormer | RS | 0.774 | 0.798 | 0.605 |
| 3FC_ECFP4 | RS | 0.663 | 0.854 | 0.687 |
| ECLIPSE_SELFormer | RS | 0.785 | 0.783 | 0.605 |
| ECLIPSE_ECFP4 | RS | 0.730 | 0.823 | 0.652 |
| ECLIPSE-3FC_SELFormer | RS | 0.823 | 0.803 | 0.593 |
| ECLIPSE-3FC_ECFP4 | RS | 0.708 | 0.835 | 0.660 |
| RF_SELFormer | DCS* | 1.161 | 0.419 | 0.235 |
| RF_ECFP4 | DCS | 1.238 | 0.454 | 0.154 |
| DP_SELFormer | DCS | 1.151 | 0.438 | 0.236 |
| DP_ECFP4 | DCS | 1.152 | 0.421 | 0.251 |
| 2FC_SELFormer | DCS | 1.184 | 0.411 | 0.239 |
| 2FC_ECFP4 | DCS | 1.199 | 0.441 | 0.264 |
| ECLIPSE_SELFormer | DCS | **1.073** | **0.536** | **0.383** |
| ECLIPSE_ECFP4 | DCS | 1.167 | 0.496 | 0.336 |
| ECLIPSE-2FC_SELFormer | DCS | 1.154 | 0.510 | 0.341 |
| ECLIPSE-2FC_ECFP4 | DCS | 1.260 | 0.488 | 0.320 |
| RF_SELFormer | FDS* | 1.147 | 0.418 | 0.056 |
| RF_ECFP4 | FDS | 1.225 | 0.438 | 0.010 |
| DP_SELFormer | FDS | 1.144 | 0.374 | 0.227 |
| DP_ECFP4 | FDS | 1.172 | 0.414 | 0.275 |
| 2FC_SELFormer | FDS | 1.441 | 0.303 | 0.185 |
| 2FC_ECFP4 | FDS | 1.629 | 0.272 | 0.167 |
| ECLIPSE_SELFormer | FDS | 1.131 | **0.546** | 0.348 |
| ECLIPSE_ECFP4 | FDS | 1.195 | 0.518 | 0.330 |
| ECLIPSE-2FC_SELFormer | FDS | **1.072** | 0.539 | **0.375** |
| ECLIPSE-2FC_ECFP4 | FDS | 1.119 | 0.493 | 0.327 |

* FDS: fully-dissimilar-split, DCS: dissimilar-compound-split, and RS: random-split datasets.

**Table S4.** Test performance scores of the models in the ablation study.

| RANDOM SPLIT | Spearman (std) | MCC (std) |
|---:|---:|---:|
| Hpo-Dis-Path-Dti-Ccs-Ppi | 0.784 (0.005) | 0.604 (0.006) |
| Dis-Path-Dti-Ccs-Ppi | 0.774 (0.013) | 0.591 (0.015) |
| Path-Dti-Ccs-Ppi | 0.785 (0.005) | 0.604 (0.007) |
| Dti-Ccs-Ppi | 0.780 (0.002) | 0.596 (0.007) |
| Ccs-Ppi | 0.761 (0.011) | 0.582 (0.010) |
| **DISSIMILAR COMPOUND SPLIT** | | |
| Hpo-Dis-Path-Dti-Ccs-Ppi | 0.526 (0.013) | 0.379 (0.005) |
| Dis-Path-Dti-Ccs-Ppi | 0.519 (0.020) | 0.376 (0.017) |
| Path-Dti-Ccs-Ppi | 0.495 (0.018) | 0.360 (0.028) |
| Dti-Ccs-Ppi | 0.502 (0.009) | 0.344 (0.013) |
| Ccs-Ppi | 0.493 (0.016) | 0.340 (0.017) |
| **FULLY DISSIMILAR SPLIT** | | |
| Hpo-Dis-Path-Dti-Ccs-Ppi | 0.519 (0.025) | 0.350 (0.047) |
| Dis-Path-Dti-Ccs-Ppi | 0.450 (0.030) | 0.204 (0.034) |
| Path-Dti-Ccs-Ppi | 0.393 (0.048) | 0.258 (0.021) |
| Dti-Ccs-Ppi | 0.381 (0.043) | 0.268 (0.016) |
| Ccs-Ppi | 0.367 (0.029) | 0.300 (0.055) |

* Ppi: protein-protein interactions
Ccs: compound-compound similarities
Dti: drug (i.e., biotech and small-molecule) interactions with their target proteins
Path: pathway (i.e., KEGG and Reactome)-protein associations
Dis: disease (i.e., KEGG and EFO) associations with proteins, drugs, and pathways
Hpo: phenotype (i.e., HPO) associations with proteins, pathways, and diseases

**Table S5.** Docking scores for the three most active HER3-targeting compounds predicted by the ECLIPSE model.

| Compound | ECLIPSE (pChEMBL) | MolProphet (-log[mol/L]) | AutoDock Vina (*pKd* (kcal/mol)) |
|---|---:|---:|---:|
| Bosutinib | 6.26 | 6.08 | 6.00 (-8.17) |
| CHEMBL3891137 | 7.42 | 7.28 | 7.5 (-10.22) |
| CHEMBL3920824 | 7.42 | 7.53 | 7.7 (-10.49) |
| CHEMBL3905567 | 7.41 | 7.94 | 8.21 (-11.19) |

**Table S6. (a)** Node- and **(b)** edge-type statistics of the integrated CROssBAR KG.

**(a)**

| **Node Type (Source)** | **Size** |
|---|---|
| Compound (ChEMBL) | 422,617 |
| Protein (UniProt) | 23,419 |
| Phenotype (HPO) | 8,971 |
| Drug (DrugBank) | 5,420 |
| Disease (EFO) | 3,815 |
| Pathway (Reactome) | 3,184 |
| Disease (KEGG) | 1,879 |
| Pathway (KEGG) | 245 |
| **TOTAL** | 469,550 |

**(b)**

| **Node Pairs / Edge Type (Source)** | **Size** |
|---|---|
| Compound-Compound / is_similar_to (Chemfp) | 10,256,336 |
| Compound-Protein / targets (ChEMBL) - removed | 629,590 |
| Protein-Protein / interacts_with (IntAct) | 91,443 |
| Gene/Protein-Phenotype / is_associated_with (HPO) | 39,466 |
| Protein-Pathway / is_involved_in (Reactome) | 32,706 |
| Disease-Phenotype / is_associated_with (HPO) | 24,259 |
| Protein-Pathway / is_involved_in (KEGG) | 19,407 |
| Drug-Protein / targets (DrugBank) | 15,342 |
| Protein-Disease / is_related_to (EFO) | 7,270 |
| Protein-Disease / is_related_to (KEGG) | 5,911 |
| Disease-Pathway / modulates (KEGG) | 1,682 |
| Drug-Disease / indicates (KEGG) | 298 |
| **TOTAL** | 11,123,710 |
